# Adverse Effects of UV-Exposure on DNA Strand Displacement Reactions

**DOI:** 10.1101/2025.09.27.673911

**Authors:** Kutay Sesli, Yue Zhao, Natalie Kallish, Dominic Scalise

## Abstract

DNA strand displacement (DSD) reactions are widely used in molecular computing and nanotechnology due to their programmability and precise control over molecular interactions. However, experimental DSD systems often underperform relative to theoretical models, due in part to poorly characterized sources of reactant impurities. Here, we identify UV-shadowing (a standard technique for visualizing DNA during PAGE purification) as a previously overlooked cause of DSD circuit errors. Counterintuitively, existing purification protocols with UV-shadowing can increase impurities that disrupt circuit behavior. Specifically, we demonstrate that UV exposure (i) reduces DSD reaction percent yield and (ii) leads to a phenomenon we call “negative leak,” where high-concentration double-stranded complexes that are designed to be inert can sequester functional signal strands and thus disrupt circuit performance. Additionally, we find that UV damage is sequence-dependent, with adjacent pyrimidine-pair-rich strands being more susceptible to errors. To circumvent these errors, we introduce a simple and practical purification protocol that avoids UV exposure, and show that the errors no longer occur. Our results highlight UV-induced damage as a critical factor in DSD circuit performance, and suggest that UV-shadowing may have contributed to significant reproducibility and scaling challenges in the broader DSD literature.

## Introduction

DNA is a versatile tool for both molecular computation and programmable self-assembly. Specifically, DNA strand displacement (DSD) reactions (Fig. 1a) can execute complex information-processing tasks, including Boolean logic [1–3], neural network computation [4–6], and timed payload release [7, 8]. Unlike conventional electronics, the inputs and outputs of DSD circuits are molecules that can bind to and manipulate physical materials, with inputs including ion concentrations, temperature, and proteins, and outputs including light, electricity, enzyme activation, and drug release [9]. Additionally, techniques such as DNA origami [10] have enabled the self-assembly of nanostructures with complex geometries [11, 12]. Together, DNA circuits and self-assembly architectures enable algorithmic control over matter from the molecular [13] to the macroscopic [14] scale.

**Figure 1:**
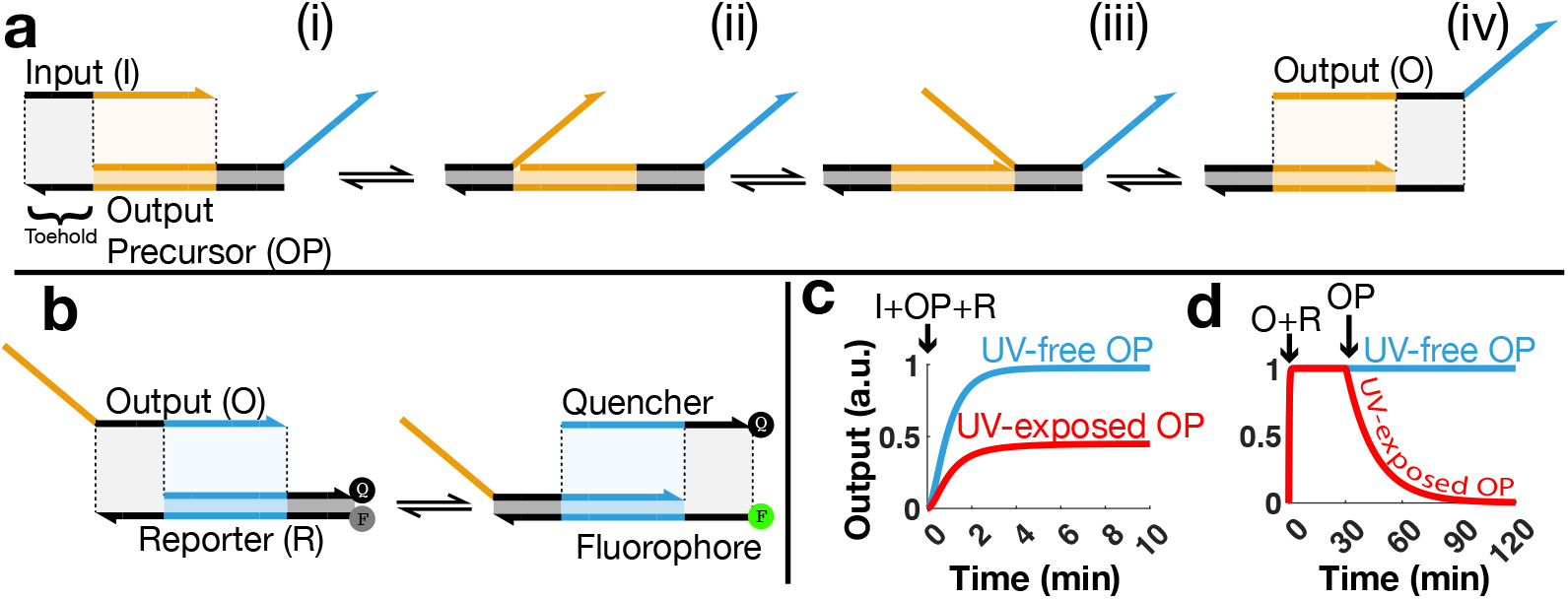
**(a)** DNA strand displacement (DSD) reaction: (i–ii) The input signal binds to its complementary overhang, called a “toehold,” on the output precursor (OP). (ii–iii) The input strand displaces the incumbent strand via the branch migration mechanism. (iii–iv) The output strand dissociates and can now participate in subsequent DSD reactions. **(b)** Reporter DSD reaction that is used to monitor the functional output signal amount in our reactions. **(c)** Percent yield drop: When the same amount of input is mixed with UV-exposed OP, a lower functional output amount is reported compared to OP not exposed to UV. **(d)** Negative leak: If the output is mixed with UV-exposed OP, especially when OP is at a higher concentration than the output, the reported output decreases.

In principle, DSD reactions are highly programmable, with the ability to implement any chemical reaction network with reaction rate constants tuned through the length of a short binding domain called a “toehold” [16] (Fig. 1a). In practice, however, experimental DSD systems often fail to match theoretical predictions. Specifically, two challenging spurious errors observed in DSD circuits include (i) failure to reach full percent yield, where the reaction yields less output concentration than theoretically expected, and (ii) “leak” reactions [17, 18] between species that are not designed to react.

In this paper, we identify **UV-shadowing during polyacrylamide gel electrophoresis (PAGE) purification as a major cause of reactant impurities that result in reduced percent yield** (Fig. 1c). We show that exposing a DNA signal strand to 3 minutes of UV-shadowing can reduce its reaction percent yield by up to ∼48%, while the same exposure on a double-stranded reversible translator complex can reduce percent yield by up to ∼74%. We explore the effects of UV on a wider parameter space including altering DNA sequence, adjusting UV exposure time, varying the UV-exposed DNA concentration in the reactions, and monitoring the reactions with reversible vs. irreversible fluorescent reporting. We further **uncover a previously unreported class of UV-induced leak reactions that arise at high reactant concentrations (relative to the signal), which we term “negative leak”** (Fig. 1d). Finally, we present a **simple purification method that avoids UV exposure** to the recovered samples, and demonstrate that this technique fully resolves the errors discussed above.

These findings suggest that many published DSD systems may have been affected by previously unrecognized UV damage, as UV-shadowing remains a commonly used method for band identification during PAGE purification in the DSD literature [3, 6–8, 17, 19–24]. Without controlling dosage, UV damage has likely contributed to difficulties in replicating DSD experiments due to randomized damage to the component strands in every experiment. Further, the discovery of UV-induced negative leak offers a new explanation for persistent leak at high concentrations. While prior studies have investigated the molecular mechanisms of UV-induced damage to nucleic acids [25] and quantified the damage severity [26–28], no studies have examined the effect of this damage on DSD reactions. Our goal in this study is to show that even brief UV exposure can significantly impair DSD circuit performance, rendering direct UV-shadowing unsuitable for purification in DSD applications. By revealing a common but correctable source of error, this work suggests new paths toward more robust and scalable DSD circuits.

## Background

### Polyacrylamide gel electrophoresis (PAGE) purification

PAGE is a standard method for purifying double-stranded complexes in DSD systems to ensure a stoichiometric 1:1 ratio of top and bottom strands and to eliminate side products that can drive unintended reactions [2–5, 7, 8, 16, 17, 19–24, 29–38]. Such purification protocols commonly include the use of UV light to visualize DNA bands [3, 6–8, 17, 19–24], because DNA absorbs UV light and creates shadows that help the experimenter locate and excise DNA bands. This gel monitoring method is referred to as UV-shadowing [26].

This widely used purification method exposes DNA to UVC light (200–290 nm), which is known to damage nucleic acids [26, 27, 39]. One of the most common types of UVC-induced damage is covalent intrastrand pyrimidine dimerization. Dimerization occurs predominantly between two adjacent thymine bases on a DNA strand, by the formation of a cyclobutane ring that covalently links them, or between adjacent cytosine–thymine, thymine-cytosine or cytosine–cytosine bases via a C6–C4 covalent linkage [27, 39]. Although UV can induce backbone breakages, at the wavelengths used for UV-shadowing, single-stranded breaks are less prevalent [40], and double-stranded breaks are unlikely to occur from direct exposure alone without enzymatic activity [39]. In principle, any intrastrand base dimerization can disrupt interstrand hybridization [39]. While UV-induced damage during PAGE is well studied for RNA [26], its impact on the kinetics and reliability of DSD circuits has not yet been systematically investigated.

### DSD reaction percent yield

We define **DSD reaction percent yield** as the amount of functional output obtained from a DSD reaction per the maximum theoretically obtainable functional output concentration. **Functional output** refers to an output DNA capable of participating in another DSD reaction with its complementary strand. For example, when running the DSD reaction shown in Fig. 1a, if we theoretically expect to measure ∼1 unit of output, but measure ∼0.45 unit, as in Fig. 1c, then the percent yield is calculated as (0.45 unit*/*1 unit) × 100 = 45%

The percent yield performance of existing DSD systems reported in the literature is often difficult to assess. A commonly used data normalization technique [1, 3, 5, 23] defines the maximum **measured** output concentration as 100% output release, even if some of the released output strands are damaged and unable to react with the reporter. This approach by definition ignores all percent yield drops, potentially masking a widespread failure to achieve reaction completion.

### Monitoring DSD reactions with reporters

DSD reaction kinetics are typically monitored by reporter reactions. A reporter is a DNA complex labeled with a fluorophore (a molecule that emits fluorescence) and a quencher (which suppresses that emission). When a signal DNA displaces the top strand of the reporter, the fluorophore is separated from the quencher and begins emitting fluorescence (Fig. 1b).

By measuring the fluorescence, we can quantify the amount of signal present in the system by calibrating fluorescence against known signal concentrations (see supplementary section 1 for details). If the interaction between the signal and the reporter is reversible, the system is called a reversible reporting system (as in Fig 1b or Fig 4a). This is particularly useful when signal concentrations change over time, as the reporter can dynamically track both signal increases and decreases. In contrast, irreversible reporter reactions (as in Fig. 5a) ideally do not report decreases in signal concentration.

## Results

To evaluate the impact of UV-exposure on DSD circuit performance, we subjected the basic components of DSD systems to varying durations of UV exposure during their PAGE purification. We then measured the amount of functional output signal released. For all experiments, we compared samples that had UV exposure during purification to control “UV-free” samples that were not exposed to UV. UV-free samples were purified using a simple new method outlined in Fig. 6. All the reactions were carried out at 25^*°*^C in standard DSD reaction buffer conditions (see supplementary section 1 for details).

### UV damaged three-way and four-way input signals

In this first set of experiments, we investigated how UV exposure affects the activity of signal strands that directly react with a reporter, which is the simplest DSD reaction. We hypothesized that UV exposure to the signals would reduce output fluorescence. Based on prior studies showing that adjacent pyrimidine pairs (TT, CT, TC, CC) are more prone to forming UV-induced intrastrand dimers [39], we further hypothesized that pyrimidine-pair-rich sequences would be more sensitive to UV damage. We tested two classes of strand displacement signals: (i) three-way DNA strand displacement (3wDSD), the most widely used DSD architecture, in which a single-stranded DNA (ssDNA) signal invades a double-stranded DNA (dsDNA) reporter, and (ii) four-way DNA strand displacement (4wDSD), a less common architecture in which a dsDNA signal interacts with a dsDNA reporter [38]. In both systems, the signal was UV-exposed for 3 minutes during purification, and we evaluated the effects of UV exposure on pyrimidine-pair-rich (PP-rich) versus pyrimidine-pair-poor (PP-poor) signal sequences to investigate sequence-dependent effects.

3wDSD experiments were performed with two types of signals differing in their sequence design alphabets: a PP-poor one with *{* A, T, G*}* and a PP-rich one with *{*A, T, C *}*, the latter of which particularly enriches the signal with pyrimidine pairs. 4wDSD experiments also used two types of signals, but this time the signals differed only in the PP-richness of the toeholds, while the dsDNA region designs remained unchanged. Both 3wDSD and 4wDSD were run at signal concentrations of 0, 12.5, 25, 37.5, and 50 nM, using both UV-exposed and UV-free signal strands, together with 75 nM of their complimentary UV-free reporters. The baseline-corrected equilibrium fluorescence values of the reporting reactions were plotted in Fig. 2. In both 3wDSD and 4wDSD experiments, UV-exposed signal reactions consistently reached lower equilibrium fluorescence values than UV-free cases, with the disparity amplified in PP-rich signals and at higher signal concentrations. The percent fluorescence drop was approximately 48% for PP-rich 3wDSD, 49% for PP-rich 4wDSD, 10% for PP-poor 3wDSD, and 11% for PP-poor 4wDSD.

**Figure 2:**
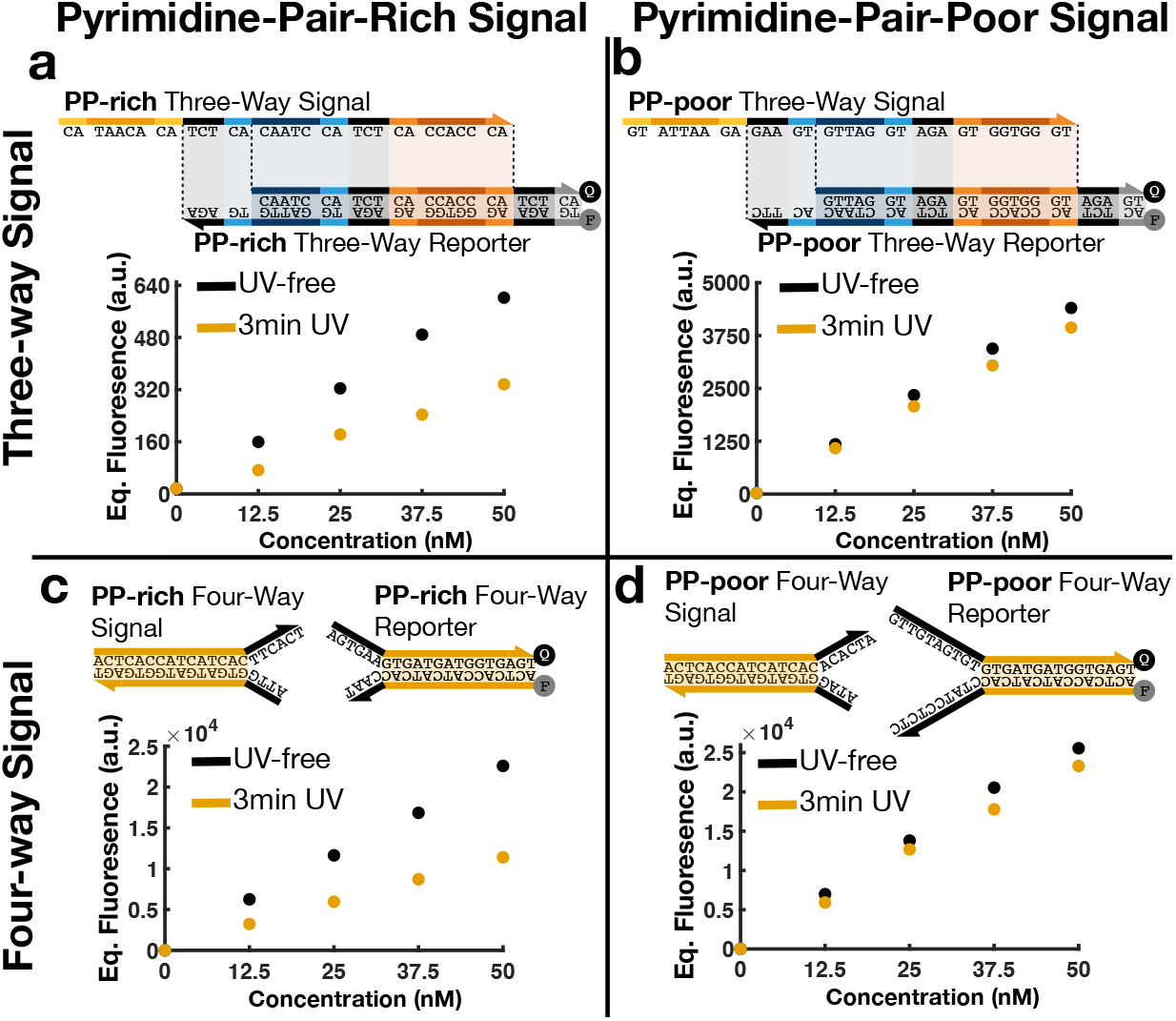
UV-shadowing reduces functional signal concentration: Equilibrium fluorescence from reporting reactions using two signal sequence types (pyrimidine-pair-rich (PP-rich) and pyrimidine-pair-poor (PP-poor)), in-house PAGE purified with and without 3-minute UV exposure, reacting with a non-in-house-purified 75 nM reporter complex. **(a)** Three-way reporting reaction with PP-rich signal. **(b)** Three-way reporting reaction with PP-poor signal. **(c)** Four-way reporting reaction with PP-rich signal. **(d)** Four-way reporting reaction with PP-poor signal.

### UV damaged dsDNA translators

Next, we explored how UV exposure affects DSD systems beyond a simple signal reacting directly with a reporter. Specifically, we tested a DSD translator reaction where an input signal reacts irreversibly with a double-stranded DNA (dsDNA) translator complex to release an output strand (see Fig. 3(a–b)). The released output strand then reacts with a fluorescent reporter complex. We hypothesized that UV exposure to the translator during purification would result in damage to some of the output strands, rendering a proportion of the output unable to react with the reporter. We investigated two scenarios for this system: one in which the reporter reaction is reversible (as in Fig. 3a), and another in which the reporter is irreversible (as in Fig. 3b). We mixed 50 nM of input signal with an excess of 70 nM dsDNA translator and a 60.5 nM reporter. This should theoretically yield fluorescence corresponding to 50 nM of output signal.

**Figure 3:**
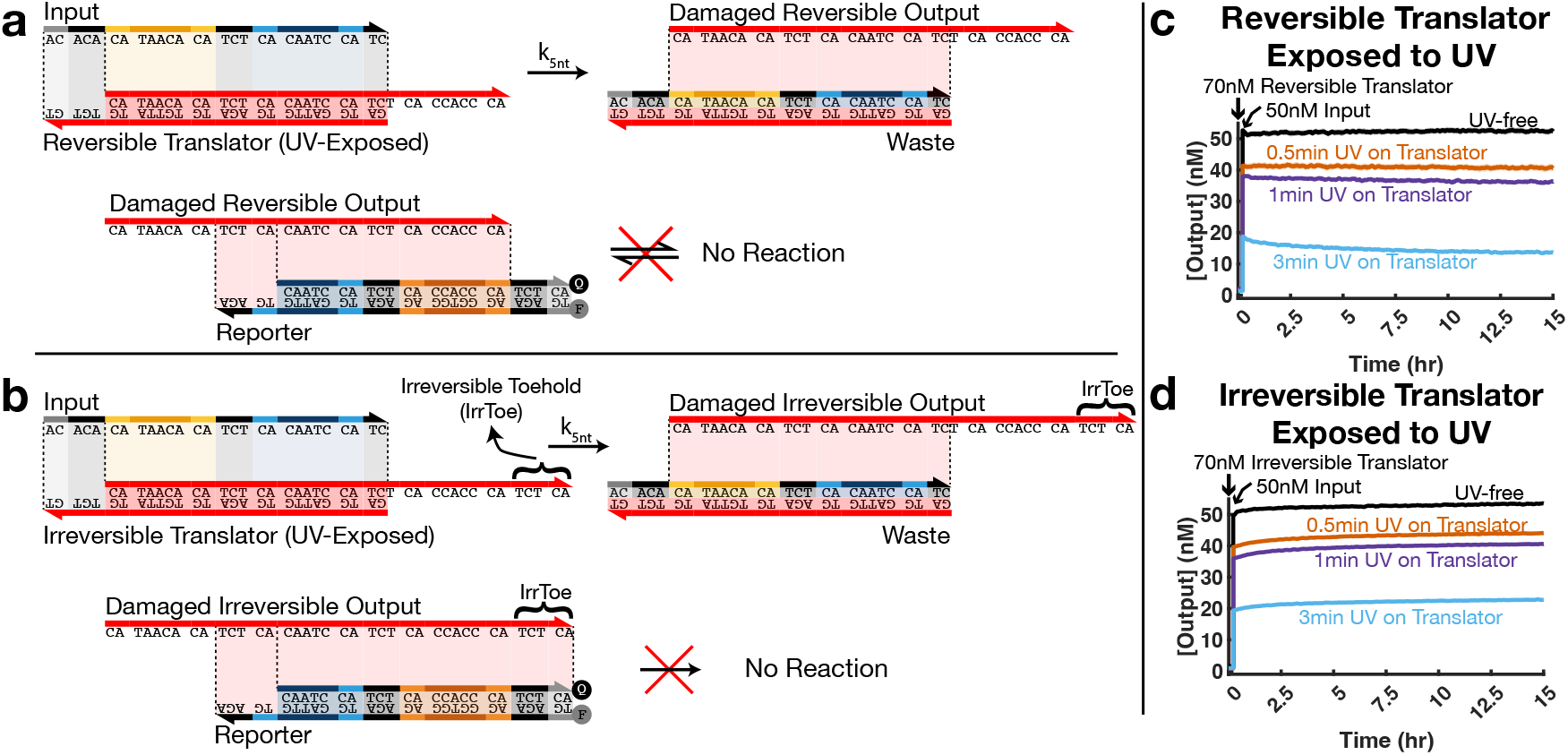
UV-shadowing reduces DNA translation reaction percent yield: **(a)** DNA translation reaction. Input signal reacts with translator to release an output that should subsequently react with the next DNA complex, which is the reporter in this case. However, when the translator is damaged by UV, the released signal cannot react with the next complex (reporter). **(b)** Reaction mechanism identical to the one in (a), except the translator releases a signal extended with a toehold that reacts with the reporter *irreversibly*. **(c)** When the translation reaction (a) is run, the output concentration measured by the reversible reporter matches the input concentration if the translator is not UV-shadowed during PAGE purification. The output signal falls below the input signal concentration if the translator is exposed to UV, exhibiting increasingly pronounced reductions at higher exposure levels. **(d)** Similarly, when translation reaction (b) is run, irreversibly reported output matches the input if the translator is not UV-shadowed during PAGE purification. The output signal falls below the input concentration if the translator is exposed to UV, exhibiting reductions that are little less pronounced than the reversible case in (c). The ideal, UV-free versions of these reactions are provided in supplementary section 6.

We observed that increasing the UV exposure time during PAGE purification caused a monotonic reduction in percent yield (Fig. 3(c–d)). Specifically, exposing the translators to 30 seconds, 1 minute, and 3 minutes of UV reduced the reported output by roughly 22%, 31%, and 74% in the reversible case, and by 18%, 24%, and 57% in the irreversible case, respectively. Under the UV-free condition, 50 nM of input DNA resulted in the release of approximately 50 nM of reported output, as expected.

These results demonstrate that even brief UV exposure during purification significantly impairs multistage DSD circuit percent yield, with the effect worsening as UV exposure increases. The effect is significant across both reversible and irreversible reporter configurations and further highlights UV-shadowing as a major source of performance loss under standard DSD conditions. The problems of UV-induced damage to dsDNA translators are particularly relevant in practice, since in-house purification is more often applied to dsDNA than to ssDNA to ensure 1:1 stoichiometry and remove undesirably formed side structures.

### UV damage generates negative leak

Finally, we uncovered a previously undocumented class of DSD leak reactions, which we call “negative leak”, that are a consequence of UV-induced damage. DSD systems are known to suffer from multiple different types of unintended leak reactions [17, 18], in which two or more species react despite not being designed to react. These leaks typically result in a gradual increase in background signal and degraded circuit specificity. Negative leak is a qualitatively different behavior in which high concentrations of UV-damaged components cause the functional signal concentration to gradually decrease over time, rather than increase.

We observe the negative leak phenomenon most clearly when examining a three-component system consisting of: (i) a single-stranded signal DNA that reacts with a reporter to generate fluorescence, (ii) a reversible reporter, and (iii) a high concentration of a signal precursor complex, in which the top strand of the precursor is an identical copy of the signal strand (Fig. 4(a–b)). In theory, the presence of the signal precursor should have no effect on fluorescence, because if a signal strand reacts with the signal precursor, it should simultaneously displace the top strand (which is a copy of the displacing signal) resulting in no net change in functional signal concentration. Alternatively, if the signal precursor complex had a leak reaction with the reporter, it should displace the top strand of the reporter to increase fluorescence.

**Figure 4:**
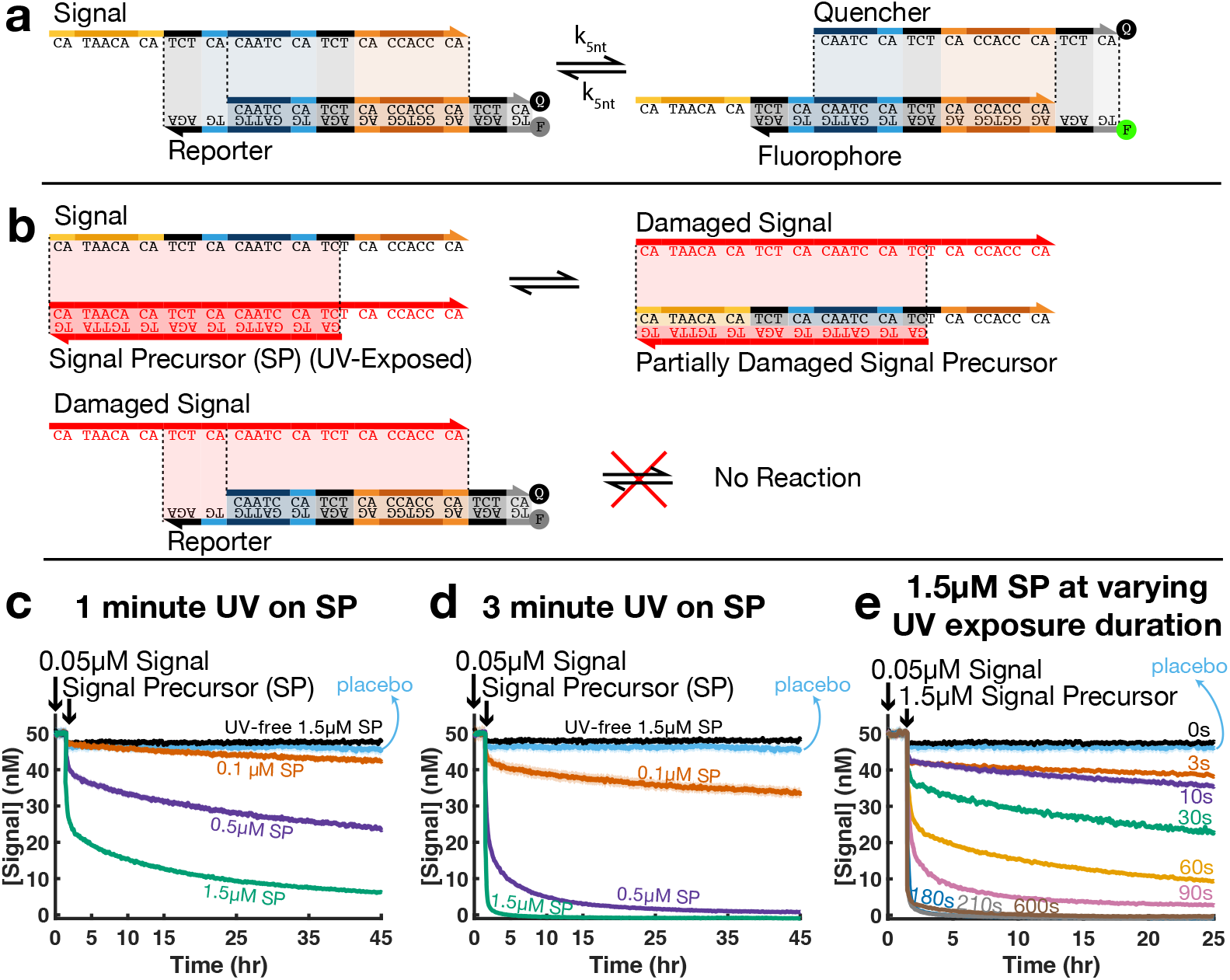
UV-shadowed DNA signal-precursors induce negative leak in reversibly reported systems: **(a)** Schematic of a reporter reaction system in which a single-stranded signal DNA reversibly reacts with a reporter, separating the fluorophore-labeled strand from the quencher-labeled strand. **(b)** Proposed mechanism for negative leak caused by UV-shadowed signal precursors (SPs). The SP is a double-stranded DNA in which the top strand is the signal strand. When a PAGE purified (with UV-shadowing) SP is added to the reporter reaction system, the functional signal strand reacts with the SP and is exchanged for a nonfunctional UV-damaged signal strand that cannot react with the reporter. This reduces the concentration of reactive signal and shifts the reporter reaction equilibrium toward the reactants. **(c)** Correlation between signal precursor concentration and the severity of negative leak for precursors UV-shadowed for 1 minute during PAGE purification. The “Placebo” data show the result when only the blank solution, containing no DNA, was added at the volume normally used when adding the signal precursor solution. **(d)** Same as (c), but using precursors exposed to UV for 3 minutes. **(e)** Correlation between UV exposure duration and the severity of negative leak for a fixed signal precursor concentration of 1.5µM.

Counterintuitively, we observed the opposite behavior: **adding UV-exposed signal precursor complex caused the reported signal concentration to decrease over time**. This result suggests that the UV-damaged signal precursor sequesters functional signal strands and, in turn, releases damaged copies of the signal incapable of reacting with the reporter (Fig. 4b), effectively removing functional signal from the system. We started our negative leak investigation by first mixing 50 nM of the reversible signal strand with the reporter. Neither the signal nor the reporter was exposed to UV, so we observed the reporting reaction to proceed as designed, as shown during the first hour and a half in Fig. 4(c–e), producing fluorescence corresponding to 50 nM. Next, we added varying concentrations of precursor, subjected to increasing durations of UV-exposure during purification. After adding the precursor, we observe that the reported signal gradually declines. The rate and extent of this decline is proportional to both the added concentration of the precursor and the duration of UV exposure. Even 3 seconds of UV exposure leads to about a 23% decline in signal within 24 hours, as seen in Fig. 4e. To measure the dilution-related concentration drop caused by the extra volume from adding the signal precursor solution, we included a placebo control in which an equal volume of blank solution containing no signal precursor DNA. The placebo showed about a 5% decline in signal due to dilution. Finally, when we added a signal precursor that was purified with no UV exposure, we observed no signal drop beyond the placebo.

We then explored the same experiment, but with an irreversible reporting reaction, in which the only difference is that the signal is extended with a toehold, rendering its reporter reaction irreversible (Fig. 5(a–b)). We observed that negative leak is still present but less pronounced with an irreversible reporter (Fig. 5(c–d)). We hypothesize that this occurs because the functional signal reacts with the reporter **irreversibly** and faster, reducing equilibrium shifts caused by subsequent consumption of remaining functional signal by the damaged signal precursor. Nonetheless, a gradual decline in the reported signal is observed over the long term, likely because the reporting reaction is not perfectly irreversible and retains a small degree of reversibility via 0-nt toehold mediated DSD in the reverse direction [16]. Notably, at low precursor concentrations, when the precursor is at 100 nM and the irreversible signal at 50 nM, the effect of negative leak is negligible for both 1-minute and 3-minute UV exposure durations. This signal-to-precursor concentration ratio (1:2) combined with an irreversible reporting system aligns with the general DSD design specifications of prior DSD computation studies [1, 3, 4, 16], which likely explains why the negative leak phenomenon has not been reported previously.

**Figure 5:**
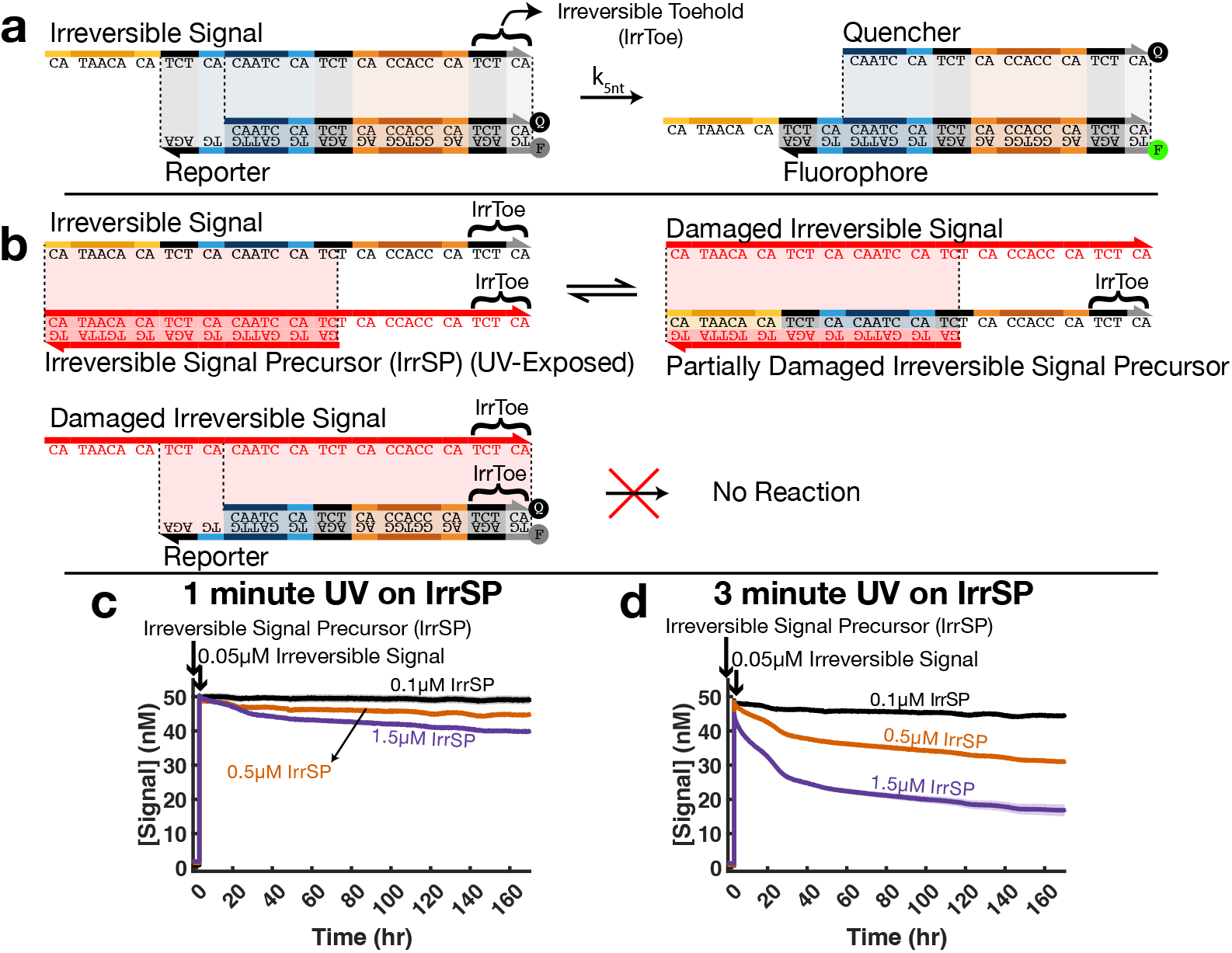
UV-shadowed DNA signal-precursors induce negative leak in irreversibly reported systems: **(a)**Schematic of a reporter reaction system, similar to Fig. 4a, but in this case the single-stranded signal DNA reacts *irreversibly* with the reporter due to its toehold extension. **(b)** Proposed mechanism for negative leak caused by UV-shadowed signal precursors. While the mechanism is identical to Fig. 4b, its impact depends strongly on its rate relative to the reporter reaction, since negative leak cannot effectively shift the reaction equilibrium of the irreversible reporting. **(c)** Correlation between irreversible signal precursor concentration and the severity of negative leak for signal precursors UV-shadowed for 1 minute during PAGE purification. **(d)** Same as (c), but with precursors exposed to UV for 3 minutes.

Based on these observations, addressing negative leak is particularly important for exploring dynamic DSD circuits, which tend to rely on reversible reactions and high precursor concentrations where negative leak is most disruptive [19].

### UV-free non-denaturing PAGE purification

For UV-free purification, we used a standard PAGE purification protocol (see supplementary section 1), with one important modification during the band visualization step. To locate the DNA band without directly exposing the final DNA sample to UV, we used a sacrificial slice of gel as explained and illustrated in Fig. 6. This method enabled visualization of the target DNA band while protecting the rest of the gel from UV damage.

**Figure 6:**
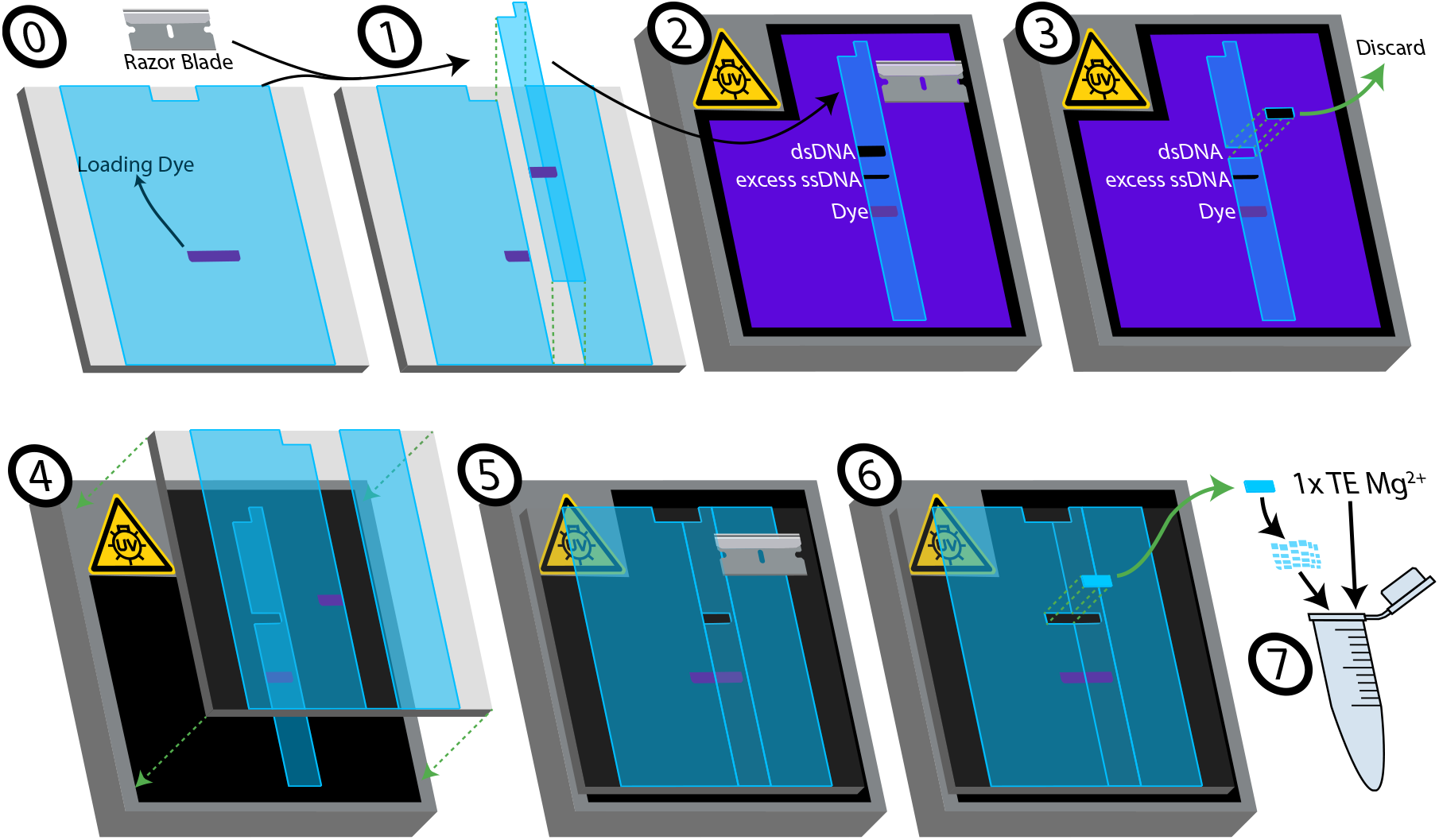
A procedure for locating DNA bands without direct UV exposure: (0) After the electrophoresis, only the loading dye band is visible to the eye. (1) A sacrificial vertical strip of the gel containing the dye band is excised with a disposable razor blade sterilized with ethanol. The strip is cut to include approximately 50% of the DNA band, though a smaller portion may also be used. (2) Only this strip is exposed to UV light to locate the dsDNA band, while the remainder of the gel remains shielded from UV. (3) The dsDNA band located in the sacrificial strip is excised and discarded. (4–5) UV light is then turned off, and the unexposed gel is aligned with the sacrificial strip at the cut-out region and placed on top. (6) By using the vertical position of the sacrificial dsDNA band and the horizontal position of the dye below, the corresponding unexposed dsDNA band is located and excised. (7) The excised gel chunk is chopped into finer pieces, placed into a 1.5 mL centrifuge tube, soaked with 1 × TE Mg^2+^ buffer, and left overnight for the remainder of the protocol (see supplementary section 1).

## Discussion

This study demonstrates that UV-shadowing, a standard technique used during polyacrylamide gel electrophoresis (PAGE) purification, introduces substantial and previously unrecognized damage to DNA that directly impairs the performance of DNA strand displacement (DSD) reactions. Despite its widespread use in the field, UV-shadowing alters the functional composition of purified strands in ways that compromise the reliability, predictability, and scalability of molecular systems based on DSD.

Our experiments show that even a few seconds of UV exposure to either single-stranded signals or double-stranded complexes can significantly disrupt DSD reaction percent yield, with the extent of signal loss increasing with longer exposure times and higher adjacent pyrimidine-pair content. Three minutes of exposure to single-stranded DNA signals resulted in up to a 48% decrease in percent yield. More severely, three minutes of exposure to double-stranded DNA translator complexes resulted in up to a 74% decrease in percent yield.

Additionally, we reported that a UV-exposed DNA signal precursor can act as a thermodynamic sink and consume the free functional signals in the reaction, a new phenomenon we refer to as “negative leak”. Negative leak is pronounced when a DNA signal is present alongside its corresponding double-stranded precursor at significantly higher concentrations and is measured using a reversible reporter. However, depending on the UV dosage, negative leak can also be observed at lower signal-to-precursor concentration ratios (e.g., 1:2) and in systems using irreversible reporters.

To circumvent these adverse effects, we presented an alternative method for locating DNA bands in a PAGE gel without direct UV-exposure. In this approach, the experimenter excises a narrow sacrificial strip from the gel that includes part of the loading well to determine the vertical position of the desired bands. Only this strip is exposed to UV, and the vertical location information it provides is then used to identify the position of the target DNA band without further UV exposure.

These findings have fundamentally important implications both within and beyond DNA strand displacement (DSD) circuits. Every paper on DSD circuits in our literature review either exposed DNA to UV during purification, or was ambiguous about how bands were imaged (and thus likely also used standard UV shadowing techniques), or did not use purified DNA (see Table S2). This suggests that essentially all published DSD research may have unknowingly suffered from reduced performance, randomized reductions in percent yield, and concentration-dependent signal loss due to UV-induced damage, potentially contributing to widespread past reproducibility and scaling challenges. Moreover, the mechanisms of UV damage described here could disrupt base-pairing interactions in any DNA-based system beyond DSD, including DNA nanostructures, aptamers, or hybridization-based biosensors. By adopting UV-free purification methods, researchers can eliminate a major but correctable source of experimental noise, improving the reliability of both DSD experiments and the broader field of DNA nanotechnology.

## Methods

All experimental methods used, including (1) DNA sequence design, (2) DNA strand acquisition and handling, (3) double-stranded complex formation, (4) non-denaturing PAGE purification, (5) DNA quantification, (6) plate reader conditions for fluorescence data, and (7) reporter fluorescence calibration and data processing, are detailed in supplementary section 1.

## Acknowledgements

This work was funded by NSF CAREER award NSF-CCF-FET-2341011 to D.S.

