## Supporting Information for "Adverse Effects of UV-Exposure on DNA Strand Displacement Reactions"

Kutay Sesli[1], Yue Zhao[1], Natalie Kallish[1], and Dominic Scalise[1,*]

[1]Washington State University, Chemical Engineering and Bioengineering, Pullman, 99163, United States

September 26, 2025

### 1. Materials and Methods

#### 1.1. DNA Sequence Design

The sequences for the four-way and PP-poor three-way experiments were designed from scratch by following the sequence design rules in [1]. Other DNA sequences were either directly taken or adapted from [2], which follow the double-long domain (DLD) motif of the leakless architecture [3, 4]. For example, our “Signal Precursor” corresponds to their ([2]) “Source,” our reversible signal to their ([2]) “X”, and our reporters are identical. We also designed additional complexes (e.g., irreversible signal-precursor, irreversible signal, and input signal) by adding extra toeholds and/or truncating domains. All sequences were analyzed using NUPACK [5, 6] to ensure that no DNA strand or reaction forms undesired secondary structures, and we used QGRS mapper [7] to confirm that our DNA strands do not form G-quadruplexes. Table S1 lists all the DNA strand sequences used in this study.

Table S1: List of DNA sequences used in this study

| Sequence Name | IDT Purification | Sequence |
|---|---|---|
| Translator Bottom | PAGE | 5’-GATGGATTGTGAGATGTGTTATGTGTGT-3’ |
| Input | PAGE | 5’-ACACACATAACACATCTCACAATCCATC-3’ |
| Irreversible Output & Irreversible Signal | PAGE | 5’-CATAACACATCTCACAATCCATCTCACCACCCATCTCA-3’ |
| Signal Precursor Bottom | PAGE | 5’-GATGGATTGTGAGATGTGTTATG-3’ |
| Revers. Output/Signal & PP-rich Three-way Signal | PAGE | 5’-CATAACACATCTCACAATCCATCTCACCACCCA-3’ |
| Signal (HPLC purified) | HPLC | 5’-CATAACACATCTCACAATCCATCTCACCACCCA-3’ |
| Quencher & PP-rich Three-way Reporter Top | HPLC | 5’-CAATCCATCTCACCACCCATCTCA/3IABkFQ/-3’ |
| Fluorophore & PP-rich Three-way Reporter Bottom | HPLC | 5’-/56-FAM/TGAGATGGGTGGTGAGATGGATTGTGAGA-3’ |
| PP-poor Three-way Signal | PAGE | 5’-GTATTAAGAGAAGTGTTAGGTAGAGTGGTGGGT-3’ |
| PP-poor Three-way Reporter Top | HPLC | 5’-GTTAGGTAGAGTGGTGGGTAGAGT/3IABkFQ/-3’ |
| PP-poor Three-way Reporter Bottom | HPLC | 5’-/56-FAM/ACTCTACCCACCACTCTACCTAACACTTC-3’ |
| PP-rich Four-way Signal Top | PAGE | 5’-ACTCACCATCATCACTTCACT-3’ |
| PP-rich Four-way Signal Bottom | PAGE | 5’-ATTGGTGATGATGGTGAGT-3’ |
| PP-rich Four-way Reporter Top | HPLC | 5’-AGTGAAGTGATGATGGTGAGT/3IABkFQ/-3’ |
| PP-rich Four-way Reporter Bottom | HPLC | 5’-/56-FAM/ACTCACCATCATCACCAAT-3’ |
| PP-poor Four-way Signal Top | PAGE | 5’-ACTCACCATCATCACACACTA-3’ |
| PP-poor Four-way Signal Bottom | PAGE | 5’-ATAGGTGATGATGGTGAGT-3’ |
| PP-poor Four-way Reporter Top | HPLC | 5’-GTTGTAGTGTGTGATGATGGTGAGT/3IABkFQ/-3’ |
| PP-poor Four-way Reporter Bottom | HPLC | 5’-/56-FAM/ACTCACCATCATCACCTATCCTCTC-3’ |
| PolyT20 | Standard Desalting | 5’-TTTTTTTTTTTTTTTTTTTT-3’ |

#### 1.2. DNA Strand Acquisition and Handling

All DNA strands were synthesized by Integrated DNA Technologies (IDT). Reporter strands were labeled with FAM as the fluorophore and Iowa Black® FQ as the quencher; all labeled strands were HPLC-purified by IDT. The remaining unlabeled strands were PAGE-purified by IDT. All DNA strands were received in dry (lyophilized) form, hydrated in 1× Tris-EDTA (TE) buffer (1× = 10 mM Tris with 1 mM EDTA) to a target concentration of 1000 µM, and quantified using the method explained in Section 1.5. The quantified strands were stored at −20 °C.

#### 1.3. Double-Stranded Complex Formation

DNA samples were annealed at concentrations of 50–100 µM in volumes of 50–100 µL, depending on the experimental requirements, with top strands in 50% excess (20% top excess for all complexes in the four-way DSD experiments) to ensure that only the purest top strands participated in hybridization; all non-four-way DSD reporters were annealed with a 120% top strand excess. Annealing was carried out by heating to 90°C, holding at 90°C for 5 minutes, and then cooling to 20°C, at a cooling rate of 1°C/min.

#### 1.4. Non-Denaturing PAGE Purification

Two-stage (5%/20%) polyacrylamide gels were prepared in dimensions of 16 cm (H) × 12.6 cm (W) × 0.80 mm (T). The bottom portion of the gel was first cast using a 20% acrylamide solution (20% acrylamide-bis 19:1 + 5.00 mM sodium borate decahydrate + 26.9 mM boric acid in Milli-Q water) up to approximately 13–15 cm from the bottom, leaving the top 1–3 cm unfilled. After polymerization of the bottom layer, the top 1–3 cm portion was cast using a 5% acrylamide solution (5% acrylamide-bis 19:1 + 5.00 mM sodium borate decahydrate + 26.9 mM boric acid in Milli-Q water). Polymerizations were initiated separately for each layer (5% and 20%) using 22 µL TEMED (Thermo Scientific) and 220 µL of 10% (w/v) APS (Ammonium persulfate) (Thermo Scientific) per 40 mL solution. The annealed DNA samples, kept at their annealing concentrations and volumes, were then mixed with *Gel Loading Dye, Purple (6X), no SDS* (New England Biolabs, B7025S) and loaded into 4 cm wide wells of the gel. Gels were run using the SE600 Standard Dual Cooled Vertical Electrophoresis Unit (Hoefer Inc.). The heat bath reservoir of the unit was filled with 1× sodium borate solution containing 5.00 mM sodium borate decahydrate and 26.9 mM boric acid in Milli-Q water. Then the heat bath was connected to a recirculating chiller (Thermo Scientific Accel 250 LC) set to 5 °C and activated at the start of the run. The heat bath was continuously stirred using a magnetic stirrer to maintain uniform temperature and ion distributions and prevent bubble formation along the edges of the gels. Electrophoresis was performed at constant 150 V until the loading dye fully passed into the 20% gel layer (taking ∼1h), then continued at 600 V for approximately 8 h. For UV-exposed species, DNA bands were visualized by UV shadowing using a UVP Benchtop UV Transilluminator: 3UV (Analytik Jena) with an 8 W bulb emitting at 254 nm, with a Kimwipe overlay. The top band (dsDNA) was excised, chopped into finer pieces, and soaked in 1× TE $Mg^{2+}$ buffer (1× TE $Mg^{2+}$ = 1× TE with 12.5 mM $Mg^{2+}$) for ∼12–24 h. Eluted DNA samples were separated from gel debris by centrifugation at 3000 rpm for 3 min, and the supernatants were collected as the final purified DNA stocks. DNA concentrations were quantified using a

NanoDrop One spectrophotometer (Thermo Fisher). If a dilution was needed for the purified stocks, they were re-quantified after dilution. For the PP-rich four-way experiment, the same procedure was followed, except the 20% gel layer was prepared with 1× Tris-acetate-EDTA (TAE) buffer (1× = 40 mM Tris, 20 mM acetic acid, 1 mM EDTA) instead of sodium borate decahydrate and boric acid.

##### 1.5. DNA Quantification

DNA quantifications were performed using a Nanodrop UV-Vis spectrophotometer. Single-stranded DNA concentrations were calculated using the following formula:

$$\text{Concentration} = \frac{A_{260}}{\varepsilon} = \frac{\text{Absorbance at 260 nm}}{\text{Extinction Coefficient}},$$

where the extinction coefficient for each strand was provided by IDT. To measure double-stranded DNA concentrations, we used the same formula, but calculated the extinction coefficient of the duplex using the following expression from [8]:

$$\varepsilon = \varepsilon_{\text{top}} + \varepsilon_{\text{bottom}} - 3200N_{\text{AT}} - 2000N_{\text{GC}},$$

where $N_{\text{AT}}$ and $N_{\text{GC}}$ are the numbers of hybridized A–T and G–C base pairs, respectively, and $\varepsilon_{\text{top}}$ and $\varepsilon_{\text{bottom}}$ are the individual extinction coefficients of the top and bottom strands comprising the duplex.

##### 1.6. Plate Reader Conditions for Fluorescence Data

Fluorescence from the reporters was captured using a BioTek Synergy H1 plate reader with the following settings: excitation wavelength, 490 nm; emission wavelength, 525 nm; optic position, bottom; gain, 75; and isothermal set-point temperature, 25°C. The experiments were conducted in Corning 284-well black clear-bottom plates. To all experimental wells, 5 µM (2 µM for the four-way experiments) PolyT20 (an oligonucleotide consisting of 20 consecutive thymine bases) was added before adding any DNA to minimize adsorption of DNA to the well walls [9].

##### 1.7. Reporter Fluorescence Calibration and Data Processing

For all experiments, reporter strands were added first, and their fluorescence values were recorded to measure the background fluorescence as a baseline. Additionally, calibration experiments were conducted alongside each experiment by adding known concentrations of signal DNA to the reporter and measuring the resulting fluorescence. The fluorescence values were baseline-corrected by subtracting their corresponding averaged baseline values (i.e., the self-fluorescence of the unactivated reporter). The correlation between known signal concentrations and baseline-corrected fluorescence was fitted to a second-order polynomial at each time point, producing a time-dependent calibration curve that inherently accounts for temporal noise.

During data processing, experimental fluorescence data were also baseline-corrected by subtracting their corresponding averaged baseline values. These baseline-corrected fluorescence values were then converted to reported signal concentration values using the time-dependent calibration curve. These processing steps can be summarized compactly by the following equation:

$$[\text{Signal}](t) = a(t) \cdot (F(t) - F_0)^2 + b(t) \cdot (F(t) - F_0) + c(t),$$

where $[\text{Signal}](t)$ is the reported signal concentration at time $t$, $F(t)$ is the measured fluorescence at time $t$, $F_0$ is the averaged baseline fluorescence, and $a(t), b(t), c(t)$ are the coefficients of the time-dependent calibration curve. Fig. S1 shows example calibration data and illustrates how a calibration curve and its coefficients were obtained for a single time point (at t = 4.00 hr in this example).

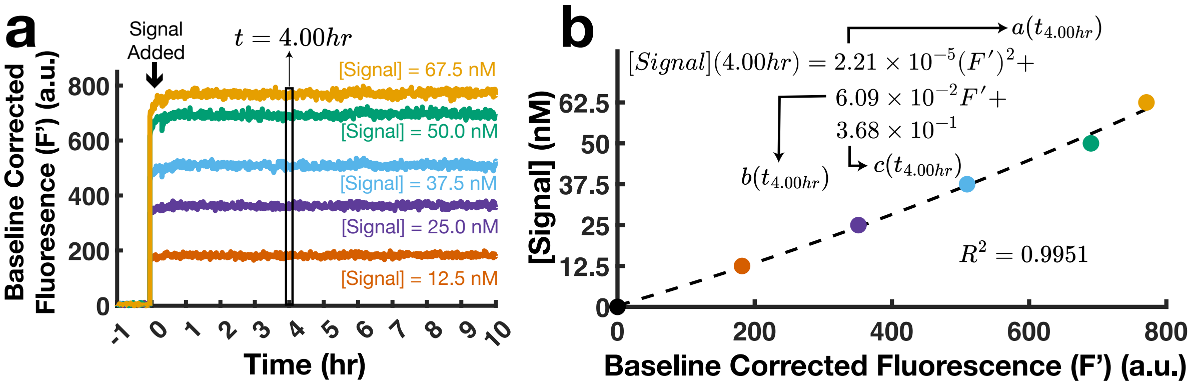

Figure S1: Illustration of an example calibration process at a single time point. **(a)** Result of a calibration curve experiment (baseline-corrected: $F' = F(t) - F_0$). **(b)** Calibration curve constructed using the data at $t = 4.00\ hr$, as also indicated in (a). The fitted coefficients were used to convert the fluorescence at $t = 4.00\ hr$ to reported signal concentration units. This procedure (fitting a curve and applying it to the fluorescence data) was repeated independently at each time point.

For experiments in which the signal DNA was added before its precursor (applicable only to Fig. 4(c–e) and Fig. S4a data), an additional rescaling step was applied. In those experiments, prior to adding the signal precursor, the reported signal occasionally deviated from their expected level (50 nM) by up to $\sim \pm 5$ nM, complicating interpretation of subsequent drops (see Fig. S2a). This deviation likely arose because the data in these figures came from independent experiments, each with its own calibration curve. Because the accuracy of the calibrations varied slightly in both positive and negative directions, directly comparing datasets calibrated independently could be misleading when comparing similar (but not identical) UV exposure time cases. Since the key feature of interest was comparing the drops in reported signals, we rescaled the datasets so that the initially reported signal before any negative leak was fixed at 50 nM. This was done by applying a two-point linear calibration on top of the prior calibration: one point fixed the initial signal to 50 nM (e.g., $54.5 \to 50$ or $45.5 \to 50$), and the second point used the control experiments (reporter + signal precursor only, with no signal) as the 0 nM reference. As a result of this rescaling, the left panel in Fig. S2a was transformed into Fig. S2b.

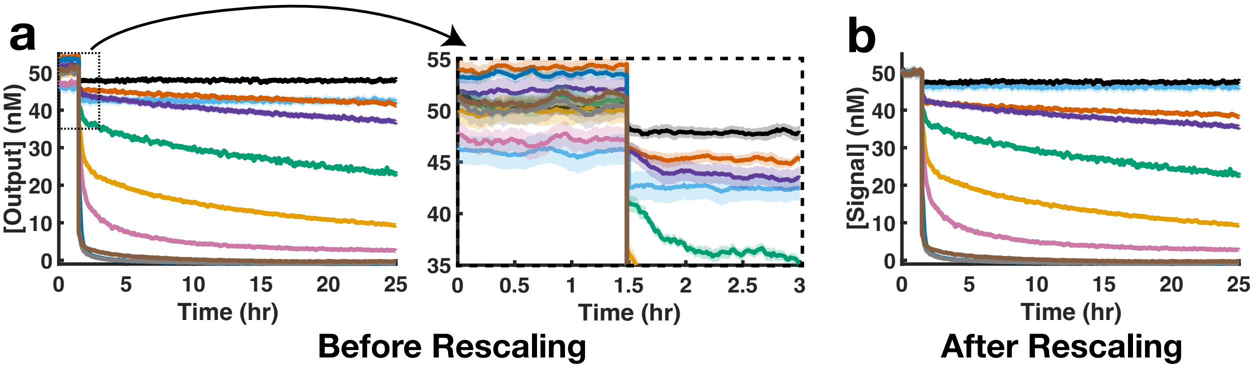

Figure S2: Illustration of the rescaling process (applied only to Fig. 4(c–e) and Fig. S4a). **(a)** Before rescaling, reported signal levels prior to signal precursor addition deviated by up to $\sim \pm 5$ nM, likely due to differences in calibration accuracy across independently run experiments. **(b)** After rescaling, the initial signal levels were aligned to 50 nM for all cases, simplifying the comparison of signal drops between similar conditions.

For those experiments that were subjected to the rescaling, the final signal expression of the data processing looks like this:

$$[\text{Signal}]_{rescaled}(t) = e \cdot [\text{Signal}](t) + f = e \cdot \left[a(t)(F(t) - F_0)^2 + b(t)(F(t) - F_0) + c(t)\right] + f$$

where $e$ and $f$ are the coefficients of the two-point linear calibration curve of the rescaling.

The experiment of each case was performed with four replicates, unless otherwise noted. One exception was the 0.1 µM IrrSP case (Fig. 5c), for which only three replicates were used because one was removed as an experimental outlier. We computed the mean of the replicates and calculated the standard error of the mean (SEM) using the following formula:

$$\text{SEM} = \frac{\text{std}}{\sqrt{n}}, \quad \text{where std} = \sqrt{\frac{1}{n-1}\sum_{i=1}^{n}(x_i - \bar{x})^2}$$

where $x_i$ represents each replicate, $\bar{x}$ is the mean of the replicates, $n$ is the number of replicates, and std is the sample standard deviation. We included the mean values of the replicates as the main data in the figures, with shaded regions representing the standard error of the mean (SEM) in both the positive and negative directions. Exceptions to this approach are Fig. 2 and Fig. S3e, where no replicates were used, and Fig. S5, where two replicates were used and their SEM was shown using conventional error bars.

Finally, we smoothed the data using a moving average with a window of 10 data points. Prior to smoothing, the dataset was segmented such that each segment began whenever new DNA (signal, signal precursor, etc.) was added. Smoothing was then applied to each segment separately to avoid the moving average from attenuating abrupt changes caused by DNA additions.

#### 2. Reported Uses of UV Shadowing in DSD Systems

We have compiled a list of papers, many of which are landmark works of great importance, that reported purifying DNA strand displacement (DSD) components for the same reasons we address: ensuring stoichiometric equality and removing poorly formed structures. Unfortunately, while these papers mention such purification, not all explicitly state whether or not UV-shadowing was used to visualize DNA bands. Instead, many use vague phrases such as "the proper bands were cut out" or "gels were imaged, and bands cut out," which are ambiguous about the use of UV. Although we strongly suspect that UV-shadowing was used in all of these cases, based on the method's popularity in the field and personal experience, we cannot confirm it with certainty. Therefore, we compiled a list of statements about the purification protocol from the relevant literature and classified each statement as either "Yes" (UV mentioned) or "Ambiguous" (see Table S2). We were not able to identify any DSD publications that did use purification after annealing, but explicitly reported that they avoided the use of UV exposure to locate the target bands.

Table S2: Reported use of DNA complex PAGE purification and UV-Shadowing in DSD Papers

| Reference | Statement | UV Mentioned |
|---|---|---|
| Fern, 2017, ACS SynBio[10] | "...the bands were cut out using UV-shadowing at 254 nm..." | Yes |
| Scalise, 2020, ACS SynBio[11] | "...cut out the purified bands using UV-shadowing at 254nm." | Yes |
| Scalise, 2018, JACS[2] | "...cut out the purified bands using UV-shadowing at 254nm." | Yes |
| Zenk, 2017, RSC Adv.[12] | "The bands... ...were identified using UV-shadowing at a wavelength of 254 nm." | Yes |
| Cherry, 2025, Nature[13] | "Target bands were cut from the gel while visualizing their shadow under UV light,..." | Yes |
| Wang, 2018, PNAS[4] | "The proper bands visualized under UV light were cut from the gels..." | Yes |
| Wang, 2023, ACS SynBio[14] | They stated that they followed the methods from this: [4] | Yes |
| Garg, 2018, Small[15] | "The gel bands were visualized under UV, excised, crushed..." | Yes |
| Eshra, 2019, IEEE TNANO[16] | "When gel finished running, DNA bands were excerpted under UV..." | Yes |
| Qian, 2011, Science[17] | "The proper bands (under UV light,...) were then cut from the gels,..." | Yes |
| Su, 2019, Nat. Commun.[18] | "The gel bands were clearly visualized under UV light, excised from the gels,..." | Yes |
| Zhang, 2009, JACS [19] | "The proper bands were cut out..." | Ambiguous |
| Srinivas, 2017, Science[20] | "The appropriate bands were cut out..." | Ambiguous |
| Zhang, 2007, Science[9] | "The proper bands were cut out..." | Ambiguous |
| Zhang, 2008, JACS[21] | "The proper bands were cut out..." | Ambiguous |
| Qian, 2011, Nature[22] | "Polyacrylamide gel electrophoresis (PAGE) purification was employed..." | Ambiguous |
| Cherry, 2018, Nature[1] | "Double-stranded complex bands were cut from the gel, chopped into pieces..." | Ambiguous |
| Lapteva, 2022, JACS[23] | "The relevant bands were incubated in 1×TE buffer..." | Ambiguous |
| Rodriguez, 2021, ACS SynBio[24] | "Bands containing the complexes were cut out from the gel..." | Ambiguous |
| Taylor, 2021, JACS[25] | "A single desired band for each complex was cut, diced, and incubated..." | Ambiguous |
| Chen, 2013, Nat. Nanotech.[26] | "The proper bands were cut from the gel..." | Ambiguous |
| Lysne, 2023, JACS[27] | "Gels were imaged, and bands cut out..." | Ambiguous |
| Zhang, 2010, Nucl. Acids Res.[28] | "The proper bands were cut out..." | Ambiguous |
| Young, 2019, ACS SynBio[29] | "Purified material was excised from the gel" | Ambiguous |
| Dabby, 2013, Caltech thesis[30] | "The proper bands were cut out" | Ambiguous |

#### 3. Reporter Brightens Irreversibly under UV

We discovered an additional UV-related phenomenon during an attempt to purify our reporter DNA (note that none of the reporters used in this work, except in this section, were in-house purified). Within the gel, we observed that the band containing the reporter DNA visibly brightened over time under UV light. To demonstrate this effect, we ran a reporter strand alongside a fluorophore strand through a PAGE gel (under previously described conditions), with the fluorophore strand serving as a reference for the maximum fluorescence intensity. Fig. S3a shows the sequence designs of these DNA strands.

Fig. S3b, recorded using a stabilized iPhone 15 with the default camera app (without any additional adjustments), shows the reporter band's increasing fluorescence intensity over time, when visualized on the UV transilluminator. To isolate this brightening effect, we sampled the average fluorescence intensity from fixed locations over the bands in the video and plotted only those regions (Fig. S3c), displaying only the green channel of the images. The dark square between 4.5–5 minutes was caused by a gel rip, which caused the gel to crawl from both sides, displacing the bands from their original positions. We then immediately returned the gel and the bands to their original positions and stabilized them using a gel-cutting ruler, which explains the appearance of the ruler after the 6-minute mark in Fig. S3b.

Fig. S3d quantifies this brightening effect. We averaged the color content in both square snippets by separately averaging the red, green, and blue (RGB) values, and then calculated the Euclidean distance between the resulting averaged RGB triplets of the reporter and fluorophore bands at each time point. This color distance was normalized such that 1 corresponds to the maximum and 0 to the minimum possible distance. The resulting plot shows a decreasing trend, indicating that the reporter is becoming increasingly similar in color to the reference. The initial increase likely resulted from camera oversaturation upon sudden UV light exposure while switching from a pitch-dark environment, and the mid-plot disturbance corresponds to the previously mentioned gel rip.

Finally, to test the irreversibility of this brightening (i.e., to confirm that it is not a temporary effect under UV light), we extracted the reporter from the gel, diluted it to nanomolar levels (it was originally loaded onto the gel at 75 µM in 50 µL), and compared it to an unexposed reporter from the same purification and gel. As shown in Fig. S3e, when fluorescence at 525 nm was measured under visible-light excitation at 490 nm, the UV-exposed reporter exhibited a much higher background fluorescence (i.e., self-fluorescence of the reporter with quenched fluorophore) and remained at these fluorescence levels for hours. Moreover, a larger increase in fluorescence was observed upon strand displacement with 50 nM signal (reversible), suggesting a lasting change in its emission and reaction characteristics.

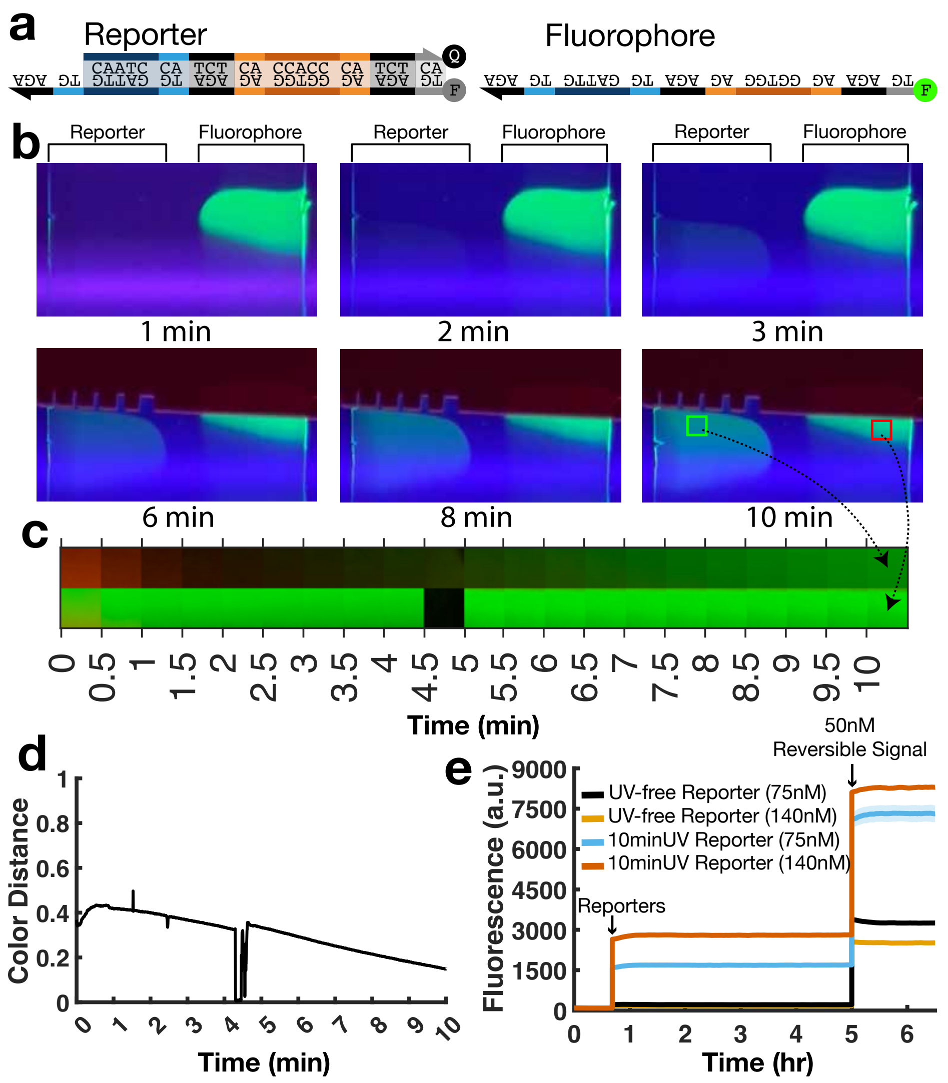

Figure S3: Effect of UV on reporter during purification **(a)** Sequence-level design of the reporter complex and the fluorophore strands, which were run side by side in separate wells through a PAGE gel. **(b)** Time-lapse gel images recorded with a stabilized iPhone 15, showing increasing fluorescence of the reporter band under continuous UV exposure. **(c)** Snippets from fixed regions of the reporter and fluorophore bands, displaying only the green color channel to highlight the brightening effect in a more isolated manner. (The dark region between 4.5–5 minutes is due to a sudden gel rip, after which the gel was returned to its original position and stabilized.) **(d)** Quantification of the visual similarity between the reporter and reference bands, calculated by taking the Euclidean color distance between the average color content of each square snippet. The decreasing trend reflects increasing similarity; early and mid-time disturbances stem from initial camera oversaturation and the gel rip, respectively. **(e)** Evidence that this brightening effect is not temporary under UV light, as shown by fluorescence measurements (525 nm) of extracted reporters using a plate reader with visible-light excitation (490 nm). The UV-exposed reporter exhibits elevated background fluorescence and a greater response to 50 nM signal (reversible), indicating a permanent alteration in both its emission and reaction behavior.

#### 4. Purifying with Fresh Gel Did Not Cause Negative Leak

It was reported in [31] that APS (ammonium persulfate), used to initiate gel polymerization, can damage RNA during PAGE purification even without UV exposure. They further observed that this effect was more pronounced when gels were used shortly after polymerization (within 1.5 hours, in their case). To test whether a similar effect occurs with DNA under our conditions, we compared freshly cast gels with older gels to assess any reduction in reported signal and whether such an effect resembles the negative leak observed after UV exposure.

To test this, we first cast and polymerized a gel and stored it at 4 °C overnight (approximately 12 hours). The next day, we prepared a second gel freshly. We then used both the fresh gel (in 1 hour of the initiation of the polymerization) and the 12-hour-old gel to purify the signal precursor shown in Fig. 4, without UV exposure. After purification, we repeated the same negative leak test presented in Fig. 4.

As shown in Fig. S4a, neither condition resulted in negative leak or signal drops. Both behaved similarly to the placebo test, in which only 1× TE $Mg^{2+}$ buffer was added as a blank solution. We also repeated this experiment by switching the DNA addition order (Fig. S4b), by adding the signal precursor first and then the signal. Under this condition, we again observed no difference between the 12-hour old and 1-hour old gels.

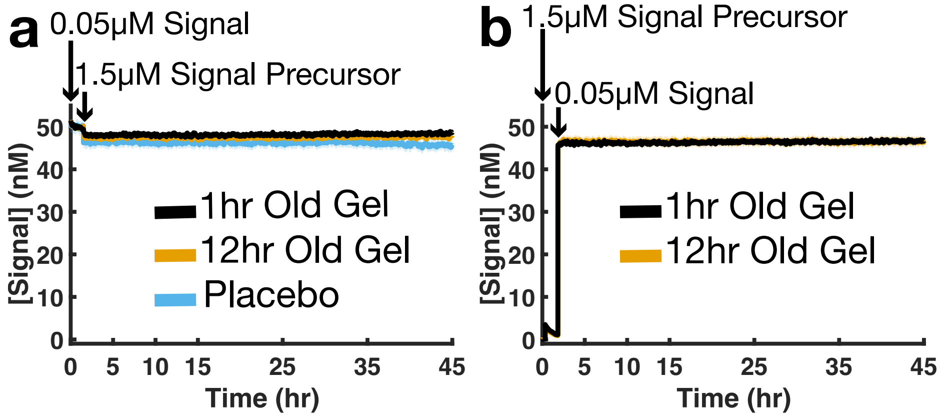

Figure S4: Fresh (1 hr old) and 12 hr-old gels (both polymerized with APS) were used to purify the signal precursor without UV exposure. **(a)** When 1.5 µM signal precursor was added on top of 0.050 µM signal, both gels behaved identically, showed no negative leak, and matched the placebo case in which only blank buffer solution was added instead of signal precursor. **(b)** Reversing the DNA addition order (adding the signal precursor first, and the signal next) also showed no difference between conditions.

#### 5. Indistinguishable Effectiveness: IDT PAGE- vs HPLC-Purified Signal DNA

We examined potential differences between strands purified by IDT using PAGE versus HPLC. Such differences could arise from how DNA bands are visualized during purification (involving UV exposure) or from variations in purification performance. When we contacted IDT regarding their use of UV during PAGE, they responded that UV is indeed used for PAGE purification, but their workflows are proprietary. To test whether these factors affect strand activity, we ran a reporting reaction (Fig. S5a) using PAGE- and HPLC-purified signal strands supplied directly by IDT, without any additional in-house purification. The same unpurified reporter (prepared as described previously) was used across all conditions to assess potential differences in functional signal concentration (i.e., the fraction of strands capable of reacting with the reporter).

Signal strands were tested at 0, 20, 40, 60, and 80 nM in the presence of 60.5 nM reporter, and the average equilibrium fluorescence was measured. This was performed for both PAGE- and HPLC-purified signal strands, with two replicates per condition. As shown in Fig. S5b, the results indicate that, within the rigor and scale of this experiment, there is no measurable distinction in the functional concentrations of IDT's PAGE- and HPLC-purified signal strands.

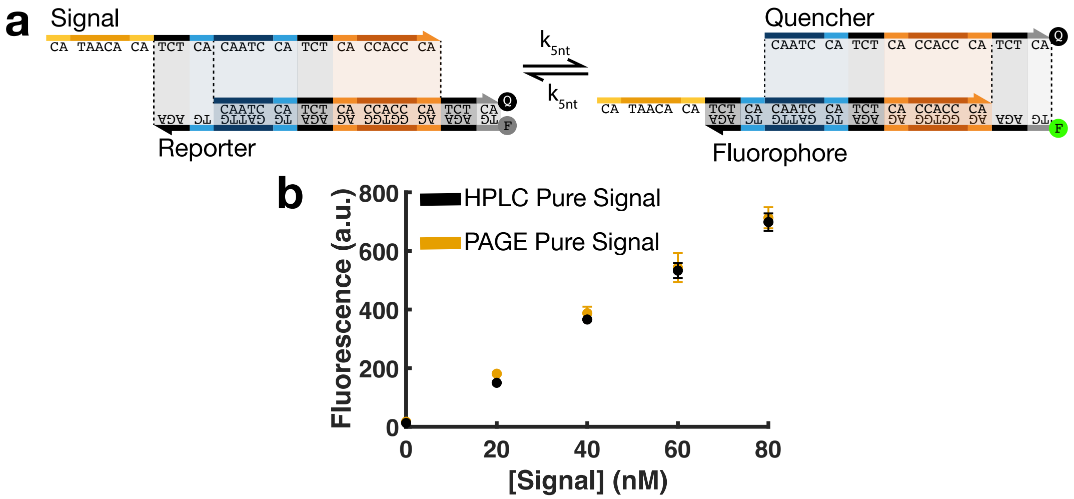

Figure S5: Comparison of IDT PAGE- and HPLC-purified signal strands. **(a)** Sequence-level design of the reporting reaction used in this experiment. **(b)** Average equilibrium fluorescence from reporting reactions containing 60.5 nM of unpurified reporter and varying concentrations (0, 20, 40, 60, and 80 nM) of signal strands purified by IDT using either PAGE or HPLC. Two replicates were performed per condition. Error bars represent the standard error of the mean. No measurable difference in functional signal concentration was observed between the two purification methods under the conditions tested.

#### 6. Ideal Reaction Network Version of Fig. 3(a–b)

This section presents how a translation DNA strand displacement (DSD) reaction would proceed under ideal conditions (see Fig. S6), as opposed to the mechanisms proposed in Fig. 3(a–b), which depicted how the translation reactions would behave if the translators were exposed to UV.

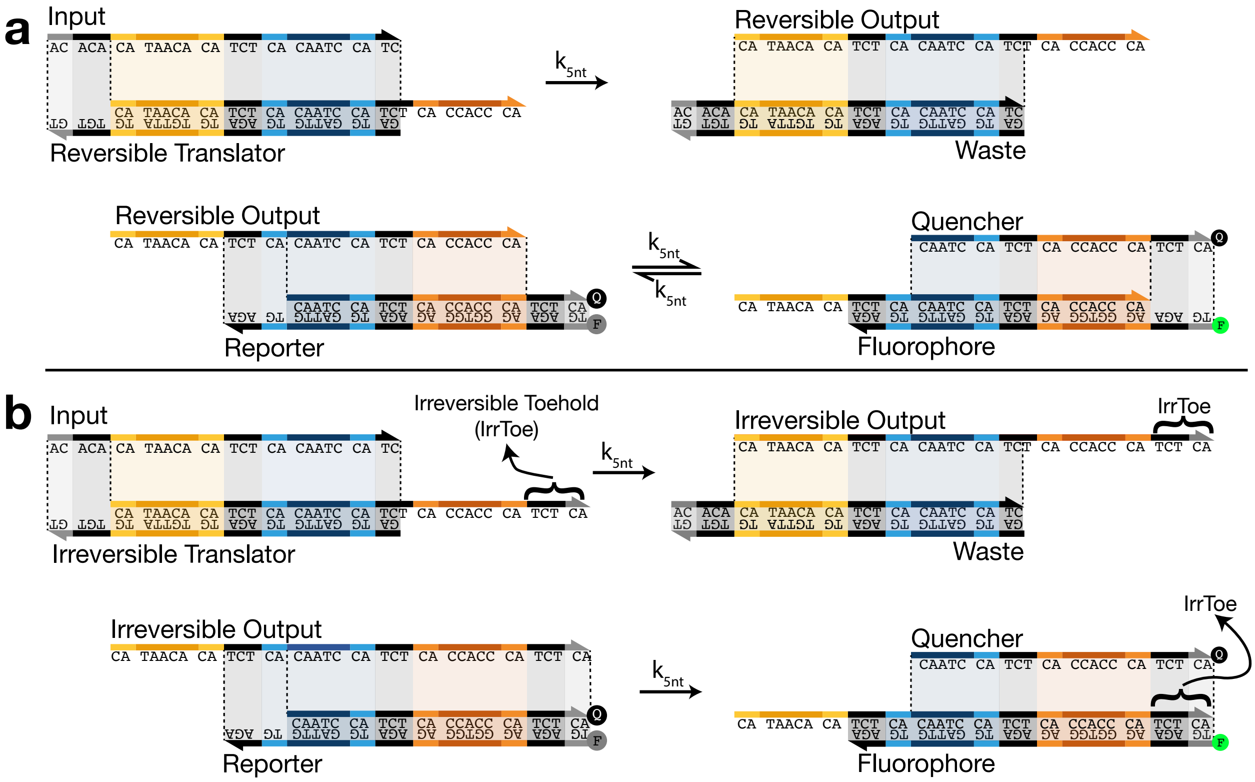

Figure S6: Ideal versions of the reactions shown in Fig. 3(a–b), with the translators free of UV-shadowing-related damage. **(a)** The input strand reacts with the reversible translator through an irreversible 5-nt toehold-mediated DSD reaction to produce a reversible output strand. This output then reacts with the reporter through a reversible 5-nt toehold-mediated DSD reaction, liberating the fluorophore from the quencher, resulting in fluorescence emission that is detected by our plate reader. **(b)** The same reaction as in (a), except the output strand reacts with the reporter irreversibly.

[7] Oleg Kikin, Lawrence D'Antonio, and Paramjeet S Bagga. Qgrs mapper: a web-based server for predicting g-quadruplexes in nucleotide sequences. *Nucleic acids research*, 34(suppl_2):W676–W682, 2006.

[30] Nadine L Dabby. *Synthetic molecular machines for active self-assembly: prototype algorithms, designs, and experimental study.* California Institute of Technology, 2013.

[31] Wipapat Kladwang, Justine Hum, and Rhiju Das. Ultraviolet shadowing of rna can cause significant chemical damage in seconds. *Scientific reports*, 2(1):517, 2012.
